# Adolescent girls at familial risk for depression with more advanced adrenarche have altered gut microbiota

**DOI:** 10.1101/2025.08.07.669149

**Authors:** A. Kolobaric, C. D. Ladouceur, A. M. Morris, B. A. Methé, E. Jašarević, L. M. Bylsma

## Abstract

**Objectives:** Rates of adolescent depression are rising, especially among girls, with children of depressed parents facing a three times higher risk. Emerging evidence suggests a link between gut microbiota, neural function, and depression risk, possibly through pathways that involve brain-body interactions, including the vagus nerve. During adolescence, sex-specific changes in the microbiota align with pubertal development, although their connection to depression vulnerability remains unclear. We compared gut microbiota in adolescents at high and low familial risk for depression and explored whether differences are affected by vagal activity and pubertal stage.

**Methods:** We collected clinical assessments, physiological data, and stool samples from 52 adolescents (aged 9-15, including 31 females), consisting of 27 high-risk and 25 low-risk individuals. We used 16S rRNA marker gene sequencing to analyze the diversity, structure, composition, and predicted function of the microbial community. A laboratory stressor task was employed to examine changes in vagally mediated heart rate variability (stress reactivity). Regressions were used to assess the relationship between depression risk, gut microbiota, and cardiovascular stress reactivity indices. Exploratory analyses investigated the effects of sex, age, and pubertal stage (adrenarche and gonarche).

**Results:** High-risk adolescents exhibited a distinct gut microbiota profile compared to low-risk adolescents, with this effect primarily driven by female participants. This profile was characterized by a higher abundance of *Prevotella*, which was 2-fold higher in high-risk females, and lower levels of other beneficial genera. High-risk females were also significantly more advanced in adrenarcheal development; the link between depression risk and adrenarcheal development was mediated by gut microbiota in females. Cardiovascular stress reactivity did not differ between groups and was not linked to gut microbiota.

**Conclusions:** Our results reveal sex-specific links between depression risk, adrenarcheal development, and gut microbiota in adolescence. The increase of Prevotella in high-risk females suggests inflammation-related pathways may connect familial vulnerability to mood disorders. Future long-term studies examining hormones, microbiota, and mood during pubertal changes are essential to determine causality and develop targeted treatments.

## Introduction

Depression is a leading cause of disability worldwide (Mulders et al., 2015; Tadayonnejad and Ajilore, 2014). Despite advances in detection and treatment, the incidence and prevalence of the condition continue to rise, especially among adolescents (Goodwin et al., 2022; Miller and Campo, 2021; Shorey et al., 2022). This increase is particularly notable among female adolescents, who are twice as likely to be affected compared to their male peers (Kalin, 2021; Kessler et al., 2001). Additionally, children whose parents have a history of depression (high familial risk) are three times more likely to develop depression themselves (Weissman et al., 2006). Many of these children also show a poor response to either pharmacological or psychotherapy interventions (Cuijpers et al., 2023; Murphy et al., 2021).

Adolescence is a developmental stage when many organ systems experience significant reorganization, including the neural circuits that regulate mood (Larsen and Luna, 2018; Paus et al., 2008; Spear, 2000). Indeed, this is when sex differences in depression rates first become noticeable, emphasizing the importance of studying sex-specific biological processes during adolescence (Breslau et al., 2017; Kessler, 2003; Nolen-Hoeksema and Girgus, 1994). While the brain has been the primary focus of research during this period, understanding how other systems and brain-body interactions change during this period could uncover new treatment targets (Pfeifer and Allen, 2021; Sisk and Zehr, 2005). One system that has received little attention is the intestinal microbiota, the community of microorganisms that resides in the gastrointestinal tract (Bastiaanssen et al., 2020; Clarke et al., 2014; Neuman et al., 2015). The microbiota contributes to host metabolism, immune regulation, and endocrine signaling, processes that also change during adolescence (Martin et al., 2019; Rastelli et al., 2019; Yang and Cong, 2021). Microbial communities go through several stages of assembly early in life, but clear sex-specific patterns only appear during puberty (Jašarević et al., 2016; Org et al., 2016; Yuan et al., 2020a). Rising gonadal hormones during adolescence drive differences in microbial composition, structure, and function between girls and boys (Korpela et al., 2021). These hormonal changes promote bacterial species that produce beta-glucuronidase enzymes involved in estrogen metabolism, creating a feedback loop where hormones influence microbial activity and the microbiome, in turn, affects circulating hormones (Baker et al., 2017; Hu et al., 2023; Li et al., 2023; Maeng and Beumer, 2023). Supporting evidence includes an increase in hormone-metabolizing bacteria in girls with precocious puberty and links between early cephalosporin exposure and earlier pubertal onset (Calcaterra et al., 2022; Hu et al., 2022). These findings suggest that environmental factors disrupting hormone-microbe interactions during adolescence could have long-lasting effects (McVey Neufeld et al., 2024; Thapa et al., 2021), prompting further investigation into microbiome connections with adolescent depression.

Considering these changes in the microbiota during adolescence, it is important to understand their possible link to depression risk. In adults, depression is linked to variations in community diversity and shifts in gut microbiota composition (J.-J. Chen et al., 2018; Chung et al., 2019; Huang et al., 2018; Kelly et al., 2016; Kolobaric et al., 2024; Lin et al., 2017; Zheng et al., 2016). Several studies have identified particular bacterial taxa and gene functions that vary between individuals with and without depression (Aizawa et al., 2016; Chahwan et al., 2019; J.-J. Chen et al., 2018; Z. Chen et al., 2018; Donoso et al., 2020; Huang et al., 2018; Jiang et al., 2015; Mason et al., 2020; Naseribafrouei et al., 2014; Valles-Colomer et al., 2019; Vinberg et al., 2019; Zheng et al., 2016). Transplanting fecal microbiota from depressed patients can cause depressive-like behavior in rodents (Kelly et al., 2016). In contrast, transferring microbiota from healthy donors can reduce symptoms in patients, supporting a causal but likely bidirectional relationship (Cai et al., 2019; Kelly et al., 2016; Rao et al., 2021). Gut microbial profiles also correlate with changes in neural activity and functional connectivity, highlighting the potential role of the microbiome in the onset and persistence of depression (Aswendt et al., 2021; Feng et al., 2022; Kohn et al., 2021). However, there is limited data connecting the microbiome to depression risk and symptoms during adolescence.

The vagus nerve represents a primary pathway through which gut microbiota affects brain function (Bistoletti et al., 2020; Bosi et al., 2020; Evrensel and Ceylan, 2015; Foster and McVey Neufeld, 2013; Fülling et al., 2019; Gershon and Margolis, 2021; Levinthal and Strick, 2020; MacQueen et al., 2017; O’Mahony et al., 2011; Rhee et al., 2009). The most robust and reliable non-invasive indicator of vagal activity is Respiratory Sinus Arrhythmia (RSA) (Berntson et al., 1993), a measure of parasympathetic nervous system activity that can be assessed by high-frequency heart rate variability (HF-HRV) (Thayer et al., 2012). Low resting values and context-inappropriate reactivity of this measure are transdiagnostic markers of poor self-regulation (Balzarotti et al., 2017) and have been observed in depression (Bylsma et al., 2015, 2014; Koch et al., 2019). Depressed individuals often exhibit blunted sympathetic reactivity, reflected by smaller changes in pre-ejection period (PEP) (Koch et al., 2019; Salomon et al., 2013). Experimental studies demonstrate that an intact vagus nerve is required for gut microbiota to induce depressive-like behavior in animals (Siopi et al., 2023), and vagal tone correlates with gut microbial composition in healthy adolescents (Michels et al., 2019).

To date, no study has simultaneously examined gut microbiota, vagal activity, and both familial and symptomatic depression risk in adolescents. Therefore, we investigated these factors in a community sample of adolescents. We tested three primary hypotheses. First, familial depression risk and current depressive symptoms would be associated with specific features of gut microbiota structure and predicted function. Second, participants at high familial risk would display lower cardiovascular stress reactivity indices, namely, respiratory sinus arrhythmia, pre-ejection period, and cardiac autonomic balance (Berntson et al., 2008). Third, group differences in gut microbiota would be mediated by vagal activity, as measured by these cardiovascular indices. Planned exploratory analyses also examined the moderating effects of age, sex, and pubertal stage.

## Methods

### Participants

Families were recruited from the Pittsburgh community. Advertisements focused on recruiting either parents with a history of depression or parents with no mental health history who had a child in the eligible age range. Exclusion criteria included a parental or youth history of mania or psychosis, primary medical conditions, current or recent use of medications that may affect study measures (e.g., antibiotics, steroids, beta blockers), and a youth history of developmental disorders, pervasive developmental disorder, head injury, and/or neurological disorder. The University of Pittsburgh Human Research Protection Office approved the study. Parents provided written informed consent, and adolescent participants provided written assent.

Before statistical analysis, we excluded 3 participants: 2 participants due to treatment with antibiotics within 3 months of providing their stool sample, as antibiotics impact the gut microbiota (Nel Van Zyl et al., 2022). One participant was excluded due to missing data on pubertal status. The final sample consisted of 52 participants, including 25 low-risk participants (10 male, 15 female) and 27 high-risk participants (11 male, 16 female). In the high-risk group, 10 participants (4 male, 6 female) met criteria for current major depressive disorder based on the Kiddie Schedule for Affective Disorders and Schizophrenia (K-SAD-PL DSM-5) semi-structured interviews (Kaufman et al., 1997). In the low-risk group, 1 participant met criteria for current major depressive disorder (1 female). Participants’ clinical and demographic information is reported in Table 1.

**Table 1.**
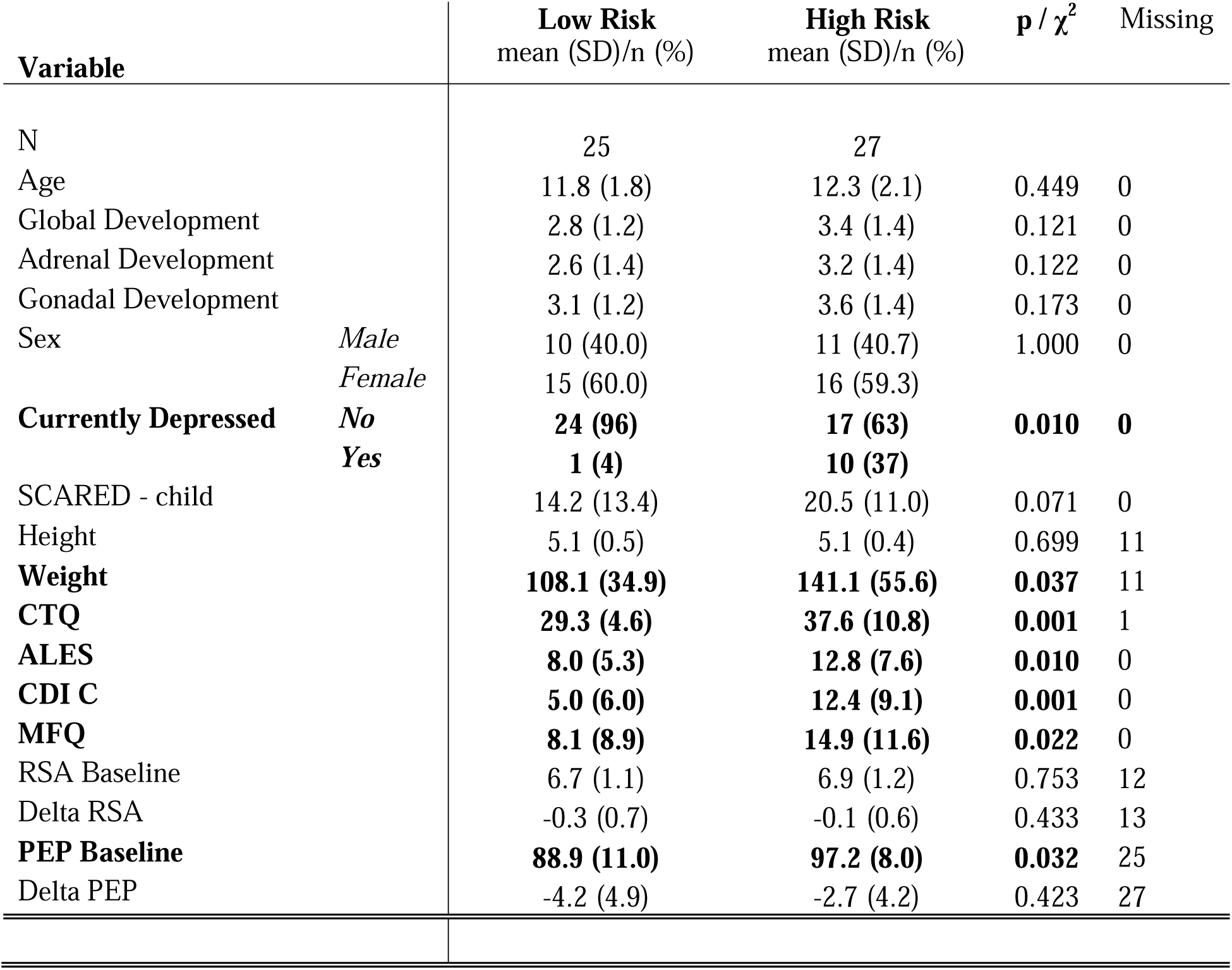
Participant sample summary. CTQ: Childhood Trauma Questionnaire. ALES: Adolescent Life Events Scale. CDI: Children’s Depression Inventory. MFQ: Mood and Feeling Questionnaire. RSA: Respiratory Sinus Arrhythmia. PEP: pre-ejection period.

### Study Design

During the first of two visits, parents and children completed a clinical diagnostic assessment and self-report questionnaires. They received their fecal collection kit, which they were asked to return before the second laboratory visit. At the second visit, typically scheduled within two weeks, the youth participated in a laboratory psychophysiological assessment.

### Self-Report Measures

Adolescents completed self-report scales of current anxiety and depressive symptoms using the Screen for Child Anxiety-related emotional disorders (SCARED) (Birmaher et al., 1999), and the Child Depression Inventory (CDI) (Kovacs, 1985), respectively. Adolescents also completed the self-report Pubertal Development Scale (PDS) (Petersen et al., 1988), which assesses five questions on physical features associated with pubertal development on a scale of 1 (no) to 4 (development seems complete). The PDS demonstrates strong alignment with physician Tanner staging and circulating hormones (Shirtcliff et al., 2009). We calculated two subscale scores representing distinct endocrine events of early adolescence: an adrenal score (average of body hair growth and skin changes items) indicating adrenarche, and a gonadal score (average of growth spurt, breast/testes development, and menarche items) indicating gonadarche. Each subscale ranges from 1 to 4, with higher scores indicating more advanced development. In addition to these subscales, we retained the traditional PDS total score and a Tanner-stage–based categorical score.

### Psychophysiological Stress Reactivity

During the second laboratory assessment, the youth completed the Trier Social Stress Task for Children (TSST-C), a well-validated measure that reliably elicits robust psychophysiological responses in youth (Kudielka et al., 2004). Electrocardiography and impedance cardiography (ECG and dZ/dt signals) were sampled continuously at 1000 Hz using a Mindware Bionex system and Cleartrace LT disposable Ag/AgCl electrodes placed in a modified Lead-II configuration. R-wave markers in the ECG signal were processed with Mindware’s MAD/MED artifact detection algorithm. Signals were manually inspected and suspected artifacts corrected using Mindware HRV and IMP Analysis software. The time series of interbeat intervals was created using an interpolation algorithm, and then linearly detrended, mean-centered, and tapered with a Hanning window. Faster Fourier analysis was used to quantify spectral power values (in ms^²^/Hz).

HF-HRV was assessed as the natural log (ln) spectral power value in the high-frequency (HF) bandwidth (0.15-0.50). Pre-ejection period (PEP) was derived from the dZ/dt signal as the time interval (ms) between ventricular contraction and the opening of the aortic valve. CAB was computed by first normalizing HF-HRV and PEP using a z-scale transformation, and PEP was multiplied by -1 to invert the relationship to a positive association (where greater PEP values indicate greater sympathetic activity). CAB was computed using the formula: CAB = HF_HRVz -(-PEPz), thus, greater CAB values indicate a greater relative balance of parasympathetic relative to sympathetic activation (Berntson et al., 2008).

### Fecal Microbiota Sample Collection

During the first laboratory session, families were given an at-home collection kit including DNA/RNA Shield Fecal Collection tubes (Zymo) for nucleic acid preservation and short-term storage at ambient temperatures. The participating parent was present during the instructions to the child so they could assist the child at home. Specimens were mailed to the University of Pittsburgh Center for Medicine and the Microbiome. Upon receipt, specimens were sub-aliquoted and then stored at -80 °C for long-term preservation.

### DNA extraction and 16S rRNA marker gene sequencing

DNA extraction was performed using the Qiagen Powersoil Microbiome Kit EP for automated DNA extraction using an Eppendorf, 5075VTC liquid handling workstation as previously described (Li et al., 2022). Extracted genomic DNA (gDNA) was amplified for the hypervariable V3V4 regions using Q5 HS High Fidelity polymerase (New England BioLabs, Ipswich, MA) with inline barcode primers design as previously described (Caporaso et al., 2012). Approximately 5-10 ng of each sample were amplified in 25 µL reactions. Cycle conditions were 98°C for 30 seconds, then 25 cycles of 98°C for 10 seconds, 57°C for 30 seconds, and 72°C for 30 seconds, with a final extension step of 72°C for 2 minutes. Amplicons were purified with AMPure XP beads (Beckman Coulter, Indianapolis, IN) at a 0.8:1 ratio (beads:DNA) to remove primer dimers, pooled and prepared as per Illumina’s recommendations (Illumina, Inc., San Diego, CA), with an added incubation at 95°C for 2 minutes immediately following the initial dilution to 20pM. The pool was then diluted to a final concentration of 7pM + 20% PhiX control (Illumina). Sequencing was done on an Illumina MiSeq 500 cycle V2 kit (Illumina).

### Statistical Analysis

All microbiota analyses were conducted following benchmarked approaches (Bastiaanssen et al., 2023a, 2023b). We used Tjazi to compute alpha diversity, which assesses the richness and evenness of microbial communities at the genus level of each participant. To examine whether familial depression risk or current depression was associated with alpha diversity, we ran a series of linear regressions with Bonferroni corrections. A typical model included the following variables: alpha diversity index ∼ depression (risk or symptoms) + age + pubertal development + sex. We calculated beta diversity, which measures dissimilarity between two groups, and then used Permutational Multivariate Analysis of Variance (PERMANOVA) with the adonis function and 1000 permutations to test for differences between the low and high-risk groups. Finally, we conducted taxonomic and predicted functional analyses to identify specific taxa and functions associated with depression risk and symptoms. We calculated a series of linear regressions between taxa and depression risk and applied the Benjamini-Hochberg Procedure at FDR<0.2 to control for False Discovery Rate, following the guidelines of previously benchmarked work^67,68^. A typical model included the following variables: genus (relative abundance) ∼ depression (risk or symptoms) + age + pubertal development + sex. Taxonomic analyses were performed on centered log-ratio transformed data aggregated at the genus level, where any unidentified taxa and taxa present in less than 80% of the participants were removed. We computed gut metabolic modules (GMMs) and gut-brain modules (GBMs) to assess whether depression risk influences predicted microbial functional pathways (Valles-Colomer et al., 2019). These modules are a set of manually curated references of metabolic pathways reported to exist in the gut microbiome (GMM) or known to be involved in gut-brain signaling in depression (GBM) (Valles-Colomer et al., 2019). GBMs and GMMs were also transformed using centered log-ratio, and we removed any modules present in less than 80% of the participants. GBMs were limited to a subset related to depression (Valles-Colomer et al., 2019). A typical model included the following variables: gut brain/metabolic module ∼ depression (risk or symptoms) + age + pubertal development + sex. We applied the Benjamini-Hochberg Procedure at FDR<0.2 to control for False Discovery Rate following the recommendations of previously benchmarked work (Bastiaanssen et al., 2023a, 2023b).

For mediation analysis, we used principal component analysis to reduce the data dimensions on genera that differed significantly between female participants with low and high familial depression risk. The first principal component served as the summary variable for the gut microbiota. Mediation was conducted using lavaan with bootstrapping (5000 resamples) to enhance the robustness of parameter estimates and standard errors (Rosseel, 2012). The model included paths for direct, indirect, and total effects. We used bootstrapping to improve the reliability of parameter estimates and standard errors. Since mediation analyses were limited to female participants (n = 31), the statistical power to detect small indirect effects was low; therefore, these results should be considered exploratory and require confirmation in larger samples.

## Results

### Familial Depression Risk and Gut Microbiota Profile

To explore the possible link between depression risk and gut microbiota, we used 16S ribosomal RNA (rRNA) marker gene profiling to determine whether microbiota characteristics vary based on group status (low-risk versus high-risk). There was a significant difference in beta diversity between low- and high-risk groups (Figure 1A), indicating that high-risk individuals had a distinctly different microbiota structure compared to low-risk individuals. Since differences in community structure can suggest changes in the relative abundance of specific microbiota, we performed a differential abundance analysis at the genus level. We found that high-risk individuals had a higher relative abundance of *Prevotella*, a genus commonly linked to increased inflammation and inflammatory disorders (Iljazovic et al., 2021). Conversely, these individuals showed lower relative abundances of *Eubacterium eligens* group, *Lachnospiraceae ND3007* group, and *Lachnospiraceae UCG 001* (Figure 1, B-E). We next examined how current depressive and anxiety symptoms relate to the gut microbiota across the entire sample and found that higher levels of current depressive symptoms were linked to a greater relative abundance of *Prevotella* (Figure 1F) and found no associations with current anxiety symptoms.

**Figure 1.**
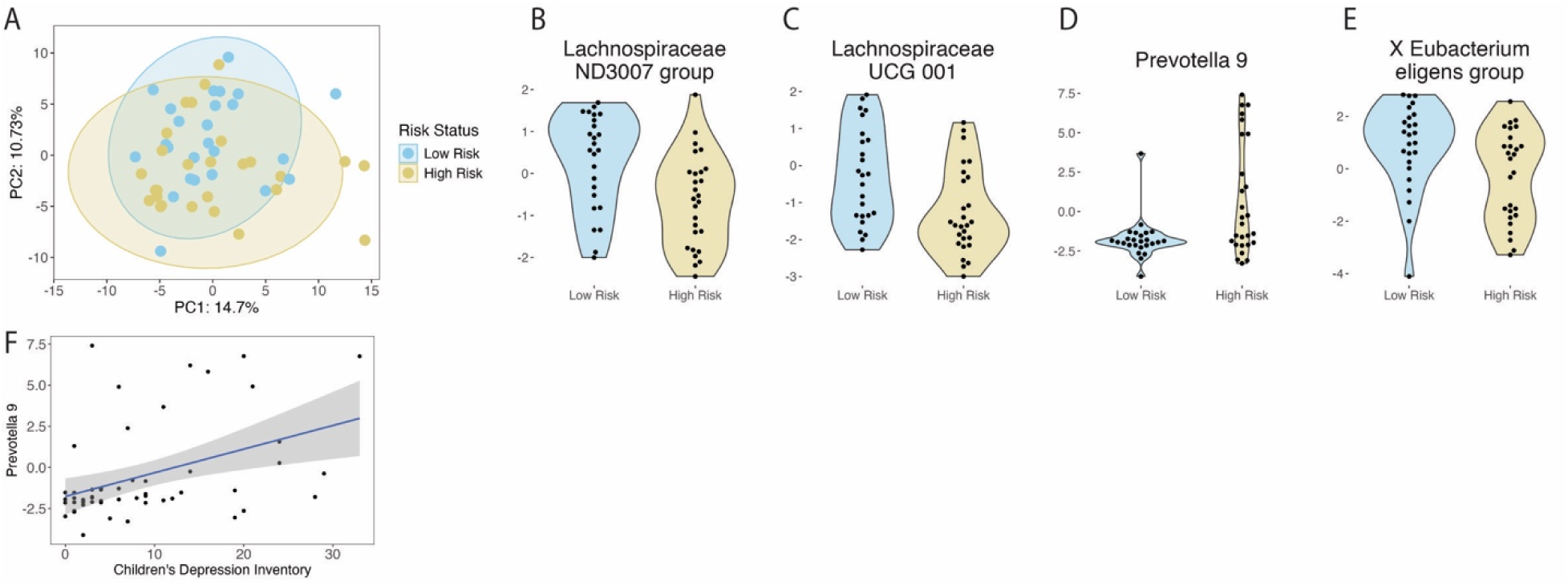
Familial Depression Risk Is Associated with a Distinct Gut Microbiota Profile. (A) Beta diversity for low and high familial depression risk groups controlling for age, sex, and pubertal development. Permutational multivariate analysis of variance (PERMANOVA) with 1000 permutations showed a significant difference in beta diversity between low and high familial depression risk groups (*F* = 1.7283, *p* = 0.0180, R² = 0.0335). (B-E) Relative abundance of genera that differ significantly between groups with low and high familial depression risk. Differences were identified using a series of regressions followed by the Benjamini-Hochberg procedure to control the false discovery rate (FDR), with a threshold of FDR < 0.2. (F) Relative abundance of genus *Prevotella* was significantly associated with current depressive symptoms, as measured by the Children’s Depression Inventory (CDI). Differences were identified using a series of regressions followed by the Benjamini-Hochberg procedure to control the false discovery rate (FDR), with a threshold of FDR < 0.2.

### Familial Depression Risk and Physiological Measures of Stress

Having observed that depression risk is significantly linked to changes in the relative abundance of key taxa, we then explored whether depression risk is connected to autonomic psychophysiology. We first assessed associations between familial depression risk and baseline measures of sympathetic activity (PEP), parasympathetic activity (HF-HRV), and cardiac autonomic balance (CAB). All analyses controlled for age, sex, and pubertal status. We found no significant associations between depression risk and any baseline physiological measures (Figure 2, A-C). In contrast, we observed significant associations between baseline HF-HRV and sex, which confirms previous findings in adults. (Figure 2D)(Snieder et al., 2007). We also found no relationship between baseline physiological measures and current depressive and anxiety symptoms across the entire sample. However, we did identify significant associations between CAB and pubertal status when examining current depressive and anxiety symptoms (Figure 2E). Next, we investigated whether familial depression risk is related to psychophysiological stress reactivity. We found no differences between depression risk groups and stress reactivity (Figure 2, F-H), nor any associations with depression or anxiety symptoms. We did, however, find associations between ΔHF-HRV and ΔCAB with age and puberty (Figure 2, I-J).

**Figure 2.**
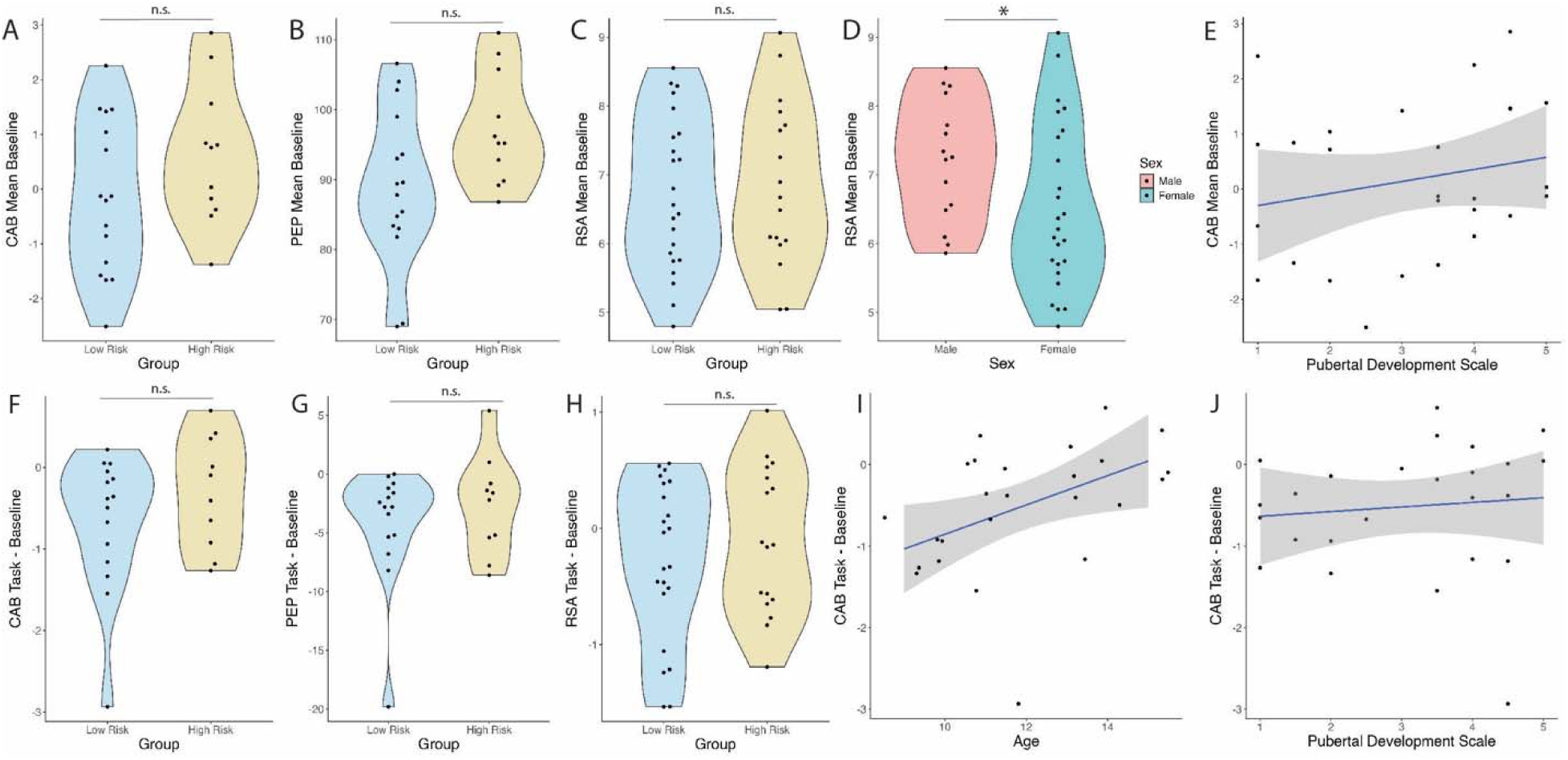
Familial Depression Risk Is Not Associated with Physiological Measures of Stress. (A-C) Baseline physiological stress measures do not significantly differ between participants with low and high familial depression risk when controlling for age, sex, and pubertal development. (D) There is a significant difference between male and female participants in measures of parasympathetic activity (RSA) when controlling for familial depression risk, age, and pubertal development. (E) There is a significant association between baseline cardiac autonomic balance (CAB) and pubertal development. F-H) Delta physiological measures of stress (calculated as Task – Baseline) do not significantly differ between participants with low and high familial depression risk when controlling for age, sex, and pubertal development. (I-J) Delta CAB is associated with Age and Pubertal Development when controlling for current depressive or anxiety symptoms and sex.

### Familial Depression Risk, Sex, and Pubertal Status

Next, we examined the relationship between depression risk and pubertal development. We found no associations between the pubertal development scale and depression risk or current depressive symptoms when controlling for age and sex (Figure 3, A-B). Pubertal development was strongly related to both age and sex. As expected, older participants showed more advanced development than younger ones, and females exhibited higher levels of development compared to males (Figure 3, C-D). Given strong evidence of sex differences in psychiatric disorders in adolescence (Campbell et al., 2021; Fox et al., 2024; Sweeting, 1995), we then stratified our sample by sex to explore sex-specific relationships between depression risk and pubertal development (Table 2, Table 3). In males, pubertal development (measured by total PDS score) was strongly associated with age, as expected, but not with depression risk or symptoms (Figure 3, E-G). Conversely, in females, pubertal development was strongly linked to familial depression risk and age (Figure 3, H-J). The PDS measures two of the three endocrine events that occur in early adolescence: adrenarche and gonadarche (Blakemore et al., 2010). Since adrenarche begins before gonadarche and they follow different developmental paths (Blakemore et al., 2010; Koopman-Verhoeff et al., 2020), we divided the PDS into two scores—the adrenarcheal and gonadarcheal development scores—to better analyze whether one or both subscales influence the relationship between pubertal development and depression risk in female participants. Using regression, we found that adrenarcheal development was strongly linked to familial depression risk (Figure 3, K-M), while gonadarcheal development was marginally associated with familial depression risk. We found no connection between depressive symptoms and pubertal status. This suggests that while pubertal development is specifically associated with depression risk, it does not relate to current depressive symptoms in adolescent females. This underscores that the risk of depression may reflect a broader, long-term vulnerability that isn’t always apparent in current symptoms within this group.

**Figure 3.**
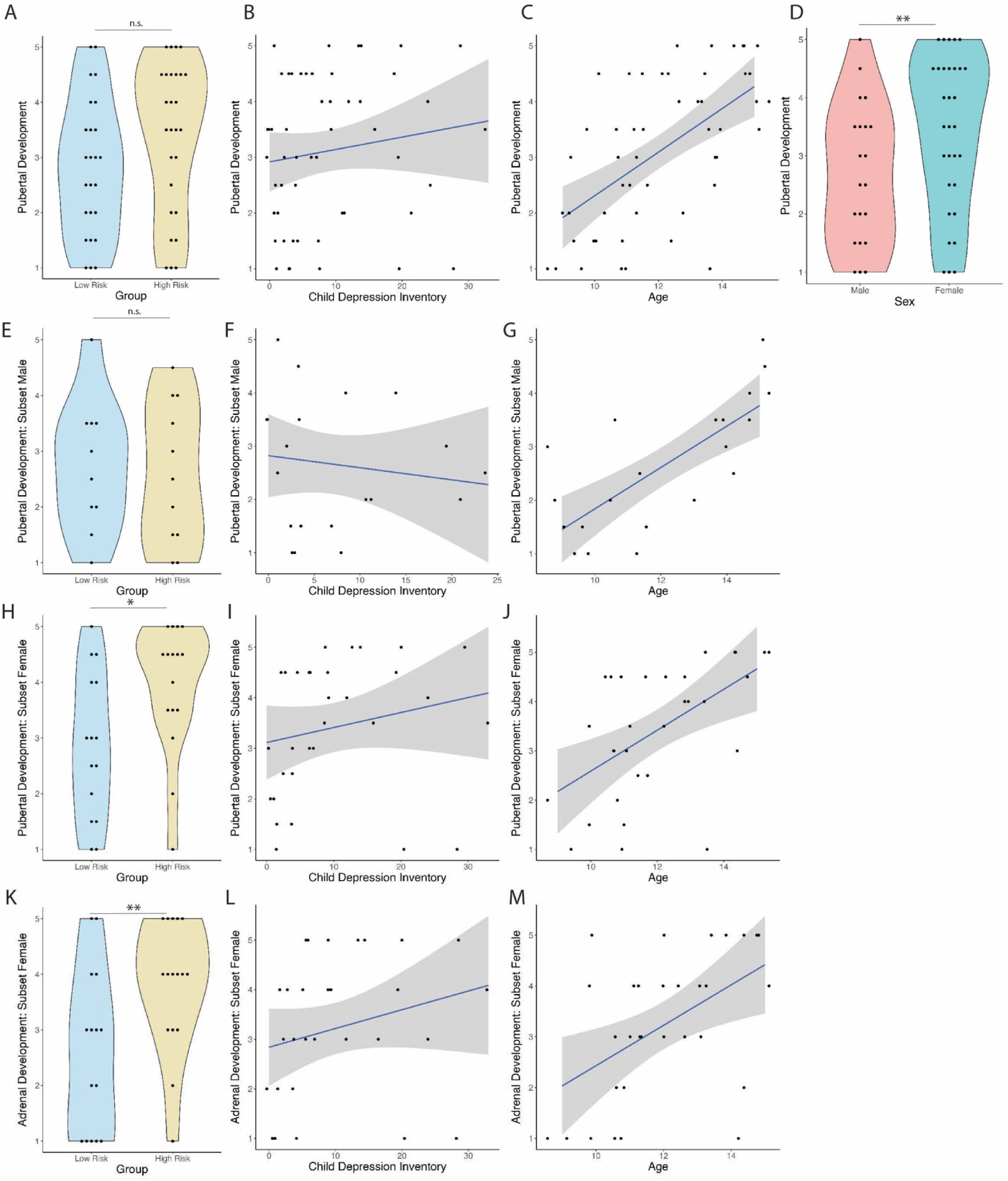
Familial Depression Risk is Associated with Pubertal Development in a Sex-Specific Manner. (A-D) Regression analysis showed that pubertal development was not significantly different between low and high familial depression risk groups (t = 1.47, p > 0.05), nor was it associated with depressive symptoms (t = -0.69, p > 0.05). However, pubertal development was significantly associated with participant age (t = 0.07, p < 0.0001) and sex (t = 2.86, p < 0.01). (E-G) In male participants, familial depression risk was not linked to pubertal development (t = -1.19, p > 0.05). Depressive symptoms also showed no association with pubertal development (t = -1.16, p > 0.05). Age, however, was related to pubertal development (t = 5.25, p < 0.0001). (H-J) In female participants, familial depression risk was associated with pubertal development (t = 2.57, p < 0.05), but depressive symptoms were not (t = -0.26, p > 0.05). Age was related to pubertal development (t = 3.51, p < 0.01). (K-M) Among female participants, familial depression risk was associated with adrenal development (t = 2.66, p < 0.05), but depressive symptoms were not (t = 0.18, p > 0.05). Age was also associated with adrenal development (t = 2.99, p < 0.01).

**Table 2.**
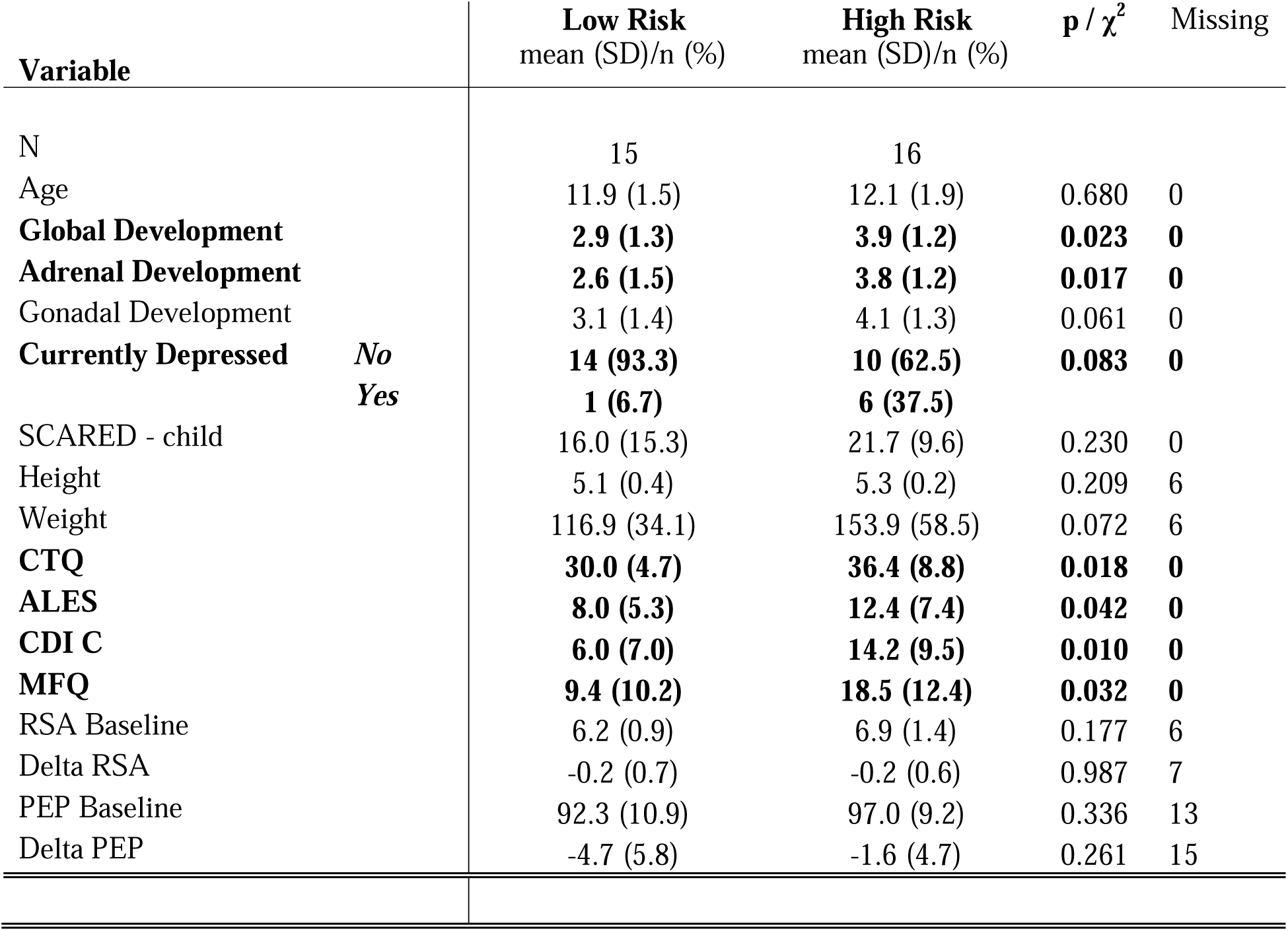
Participant sample summary, including female participants only. CTQ: Childhood Trauma Questionnaire. ALES: Adolescent Life Events Scale. CDI: Children’s Depression Inventory. MFQ: Mood and Feeling Questionnaire. RSA: Respiratory Sinus Arrhythmia. PEP: pre-ejection period.

**Table 3.** Participant sample summary, including male participants only. CTQ: Childhood Trauma Questionnaire. ALES: Adolescent Life Events Scale. CDI: Children’s Depression Inventory. MFQ: Mood and Feeling Questionnaire. RSA: Respiratory Sinus Arrhythmia. PEP: pre-ejection period.

| Variable | | Low Risk<br>mean (SD)/n (%) | High Risk<br>mean (SD)/n (%) | p / $\chi^2$ | Missing |
| --- | --- | --- | --- | --- | --- |
| N |  | 10 | 11 |  |  |
| Age |  | 11.8 (2.3) | 12.5 (2.5) | 0.540 | 0 |
| Global Development |  | 2.8 (1.2) | 2.6 (1.3) | 0.771 | 0 |
| Adrenal Development |  | 2.5 (1.5) | 2.3 (1.3) | 0.715 | 0 |
| Gonadal Development |  | 3.0 (1.1) | 2.9 (1.4) | 0.870 | 0 |
| Currently Depressed | No | 10 (100) | 7 (63.6) | 0.090 | 0 |
|  | Yes | 0 (0) | 4 (36.4) |  |  |
| SCARED - child |  | 11.4 (10.2) | 18.7 (13.1) | 0.167 | 0 |
| Height |  | 5.1 (0.6) | 4.9 (0.7) | 0.592 | 5 |
| Weight |  | 96.8 (34.4) | 115.4 (42.4) | 0.387 | 5 |
| CTQ |  | **28.1 (4.5)** | **39.5 (13.6)** | **0.022** | **1** |
| ALES |  | 8.7 (5.5) | 13.4 (8.2) | 0.139 | 0 |
| CDI C |  | **3.6 (4.1)** | **9.9 (8.3)** | **0.043** | **0** |
| MFQ |  | 6.2 (6.5) | 9.5 (8.2) | 0.313 | 0 |
| RSA Baseline |  | **7.6 (0.9)** | **6.7 (0.7)** | **0.070** | **6** |
| Delta RSA |  | -0.4 (0.7) | 0.0 (0.7) | 0.276 | 6 |
| PEP Baseline |  | **81.5 (7.3)** | **97.5 (6.7)** | **0.012** | **12** |
| Delta PEP |  | -3.0 (2.9) | -4.2 (3.5) | 0.609 | 12 |

### Familial Depression Risk, Sex, and Gut Microbiota Profile

Consistent with a sex-specific effect, we found no differences in gut microbiota profiles between low and high familial risk groups among male participants when controlling for age and adrenarcheal development. Conversely, we observed significant differences in beta diversity between low and high depression risk groups in female participants (Figure 4A). Moreover, differential abundance analysis showed that female participants at high depression risk also had higher relative abundance of *Alistipes* and *Prevotella*, as well as lower relative abundances of *Lachnospira, Lachnospiraceae ND3007 group, Lachnospiraceae UCG 001,* and *X Eubacterium xylanophilum group* (Figure 4, B-G). We also examined differences in gut metabolic modules (GMMs), which represent the gut microbiome’s potential to carry out various predicted metabolic functions (Valles-Colomer et al., 2019; Vieira-Silva et al., 2016). GMM analysis showed that female participants at high depression risk had changes in 9 gut metabolic modules, including alanine degradation II, mucin degradation, Entner-Doudoroff pathway, lactose degradation, glycerol degradation I, rhamnose degradation, galacturonate degradation I, lactose and galactose degradation, and lactaldehyde degradation (Figure 4, G-O).

**Figure 4.**
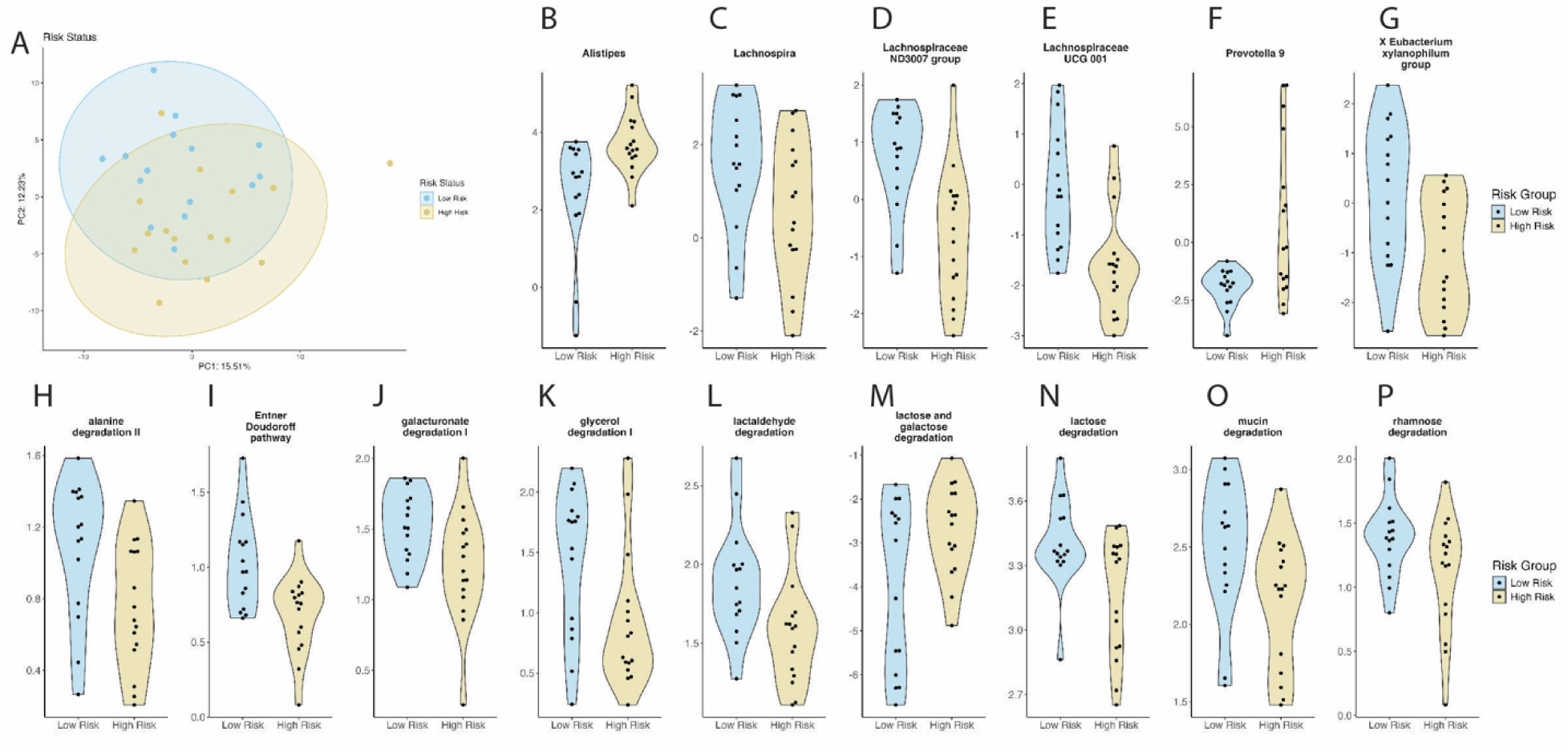
Familial Depression Risk Is Associated with Gut Microbiota in a Sex-Specific Manner. (A) Beta diversity for low and high familial depression risk groups, controlling for age and pubertal development. Permutational multivariate analysis of variance (PERMANOVA) with 1000 permutations showed a marginally significant difference in beta diversity between the low and high familial depression risk groups (F = 2.04, p < 0.01, R² = 0.065). (B-G) Relative abundance of genera that significantly differ between groups with low and high familial depression risk. Differences were identified through a series of regressions followed by the Benjamini-Hochberg procedure to control the false discovery rate (FDR), with an FDR threshold of less than 0.2. (H-P) Gut Metabolic Modules (GMMs) differ significantly between groups with low and high familial depression risk. These differences were identified through a series of regressions followed by the Benjamini-Hochberg procedure to control the false discovery rate (FDR), with a threshold of FDR < 0.2.

### Gut Microbiota Mediates the Relationship Between Familial Depression Risk and Adrenal Development in a Sex-Specific Manner

After identifying this link between family depression risk and both gut microbiota and adrenal development in female participants, we developed a hypothesis about the nature of this complex relationship. Our hypothesis is based on two main pieces of evidence. First, a recent longitudinal study showed that maternal prenatal distress accelerates adrenal development specifically in daughters (Fox et al., 2024). Second, the gut microbiota has been shown to mediate the relationship between protein intake and puberty timing in humans (Xu et al., 2024). Based on these findings, we hypothesized that the gut microbiota plays a key mediating role in connecting familial depression risk to adrenal development in female participants. Consistent with our hypothesis, a mediation model confirmed a significant direct effect of depression risk on adrenarcheal development, as well as a meaningful indirect effect mediated by gut microbiota.

During our investigation of physiological stress measures, we found that CAB was significantly linked to adrenal development across the entire sample. As a final step, we wanted to see if this relationship held for female participants only. Using regression analysis, we confirmed that baseline CAB is associated with adrenarcheal development (PDS-Adrenarche), while controlling for age, depression risk, current depressive symptoms, or current anxiety symptoms. Although our initial hypothesis that physiological stress measures mediate the relationship between familial depression risk and gut microbiota was not supported, we found that these physiological measures were strongly related to adrenarcheal development, which is influenced by both familial depression risk and the gut microbiota.

## Discussion

Our data indicate that adolescents at familial risk display a distinct gut community structure, marked by altered beta diversity and reduced abundance of genera that are typically associated with metabolic and immune benefits (Cheng et al., 2025; Leung and Weiss, 2023). Higher scores on depressive symptoms coincided with a greater relative abundance of *Prevotella*, in line with previous studies that reported the same directional association (Radjabzadeh et al., 2022; Xiong et al., 2023). While the specific causal mechanisms connecting *Prevotella* to depressive symptoms are still being uncovered, several human cohort studies have associated the expansion of this genus with systemic inflammation and mood disturbances (Larsen, 2017; Lin et al., 2023). Species within this genus produce lipopolysaccharides and capsular polysaccharides that amplify innate immune signaling, contributing to systemic low-grade inflammation often seen in patients with depressive symptoms (Larsen, 2017). *Prevotella* genomes also encode tryptophan 2,3-dioxygenase and indoleamine 2,3-dioxygenase, enzymes that redirect tryptophan from serotonin production toward kynurenine metabolites, which have neuroactive and potentially neurotoxic effects (Gao et al., 2020; Sasaki-Imamura et al., 2011; Tett et al., 2019). As this pathway has been linked to mood disorders (Zhou et al., 2023), this may point to interventions aimed at normalizing *Prevotella* that could represent a potential preventive strategy for high-risk adolescents. By contrast, we observed no gut microbial signatures linked to anxiety symptoms, a pattern that diverges from studies focusing on anxiety disorders in adults (Brushett et al., 2023; Nikolova et al., 2021; Simpson et al., 2021). This null result may be explained by the fact that our cohort consisted of individuals with low anxiety levels (i.e., only 11 participants had scores indicating a possible presence of an anxiety disorder (Runyon et al., 2018)).

Regarding vagal activity, contrary to hypotheses, we found that psychophysiological stress reactivity (HF-HRV) did not differ between depression risk groups, current depressive symptoms, or current anxiety symptoms, in contrast to prior findings (Chen et al., 2023). It is possible that we failed to detect any differences because most of our high-risk sample did not meet the criteria for current major depressive disorder. Additionally, contrary to hypotheses, vagal activity (as assessed using HF-HRV baseline levels and changes in response to a laboratory stressor) was not linked to gut microbiota. However, in both baseline and CAB, we found that when examining the relationship between depressive symptoms (but not family history of depression), pubertal status emerged as a significant predictor. This finding may suggest a complex interaction between pubertal development, stress reactivity, and familial depression risk. Interestingly, this interaction appears to be specifically related to familial risk, rather than current depressive symptoms.

Regarding sex-specific effects, analyses stratified by sex revealed that female participants mainly drove the differences in gut microbiota between low and high depression risk groups. Female participants at high risk also showed significantly more advanced pubertal development. Mediation analysis indicated that the link between depression risk and adrenarcheal development was mediated by gut microbiota in females but not in males. Additionally, in our GMM functional analyses by sex, we observed an increase in inflammation-associated genera and a decrease in genera linked to better mental health outcomes across the entire sample, predominantly driven by female participants. Furthermore, microbiota differences were evident not only at the genus level in terms of relative abundance but also in metabolic potential, covering various processes such as lipolytic fermentation, metabolism of mono- and disaccharides, mucus degradation, proteolytic fermentation, and central metabolism (Vieira-Silva et al., 2016). Building on previous work that identified sex differences in the gut microbiota across the lifespan (Jašarević et al., 2017, 2016; Markle et al., 2013), we now show that familial depression risk is associated with the gut microbiota in a sex-specific manner in adolescents.

Previous studies have highlighted sex-specific patterns of accelerated adrenarcheal development associated with adverse childhood environments and psychosocial stressors (Fox et al., 2024; Graber et al., 1995; Saxbe et al., 2015). Specifically, in a large longitudinal study, Fox and colleagues found that maternal prenatal distress predicts accelerated adrenarche, but only in girls (Fox et al., 2024). This study builds on previous findings by showing a significant link between familial depression risk and adrenarcheal development in female participants. We propose that changes in gut microbiota could serve as a potential mechanism that partly mediates the relationship between family history of depression and adrenal development. Our findings expand earlier research by identifying a modifiable mechanism that connects parental mental health with adrenal development in adolescent females. Additionally, the stage of adrenarche is linked to increased vagal activity, which makes sense given the important role of the vagus nerve in managing stress responses (Liu et al., 2021; Trevizan-Baú and McAllen, 2024).

However, establishing directionality and causality in a cross-sectional, observational sample is inherently challenging. Our mediation model was theoretically driven, based on evidence that psychosocial stressors can accelerate adrenarcheal aging in a sex-specific manner (e.g., Figure 5) (Fox et al., 2024). To examine whether gut microbiota mediates this relationship, we constructed a model with gut microbiota as a mediator between depression risk and adrenarcheal development. Notably, when we construct a model where adrenarcheal development mediates the relationship between depression risk and gut microbiota, we observe significant mediation and main effects. Our results also indicate that adrenarcheal development partially mediates the connection between depression risk and gut microbiota (Figure 6). Previous research shows that gut microbiota differs across Tanner stages in typical development (Yuan et al., 2020b). Our findings show that advanced adrenarcheal development in high-risk adolescents explains some differences in gut microbiota, but familial depression risk remains a predictor beyond this development.

**Figure 5.**
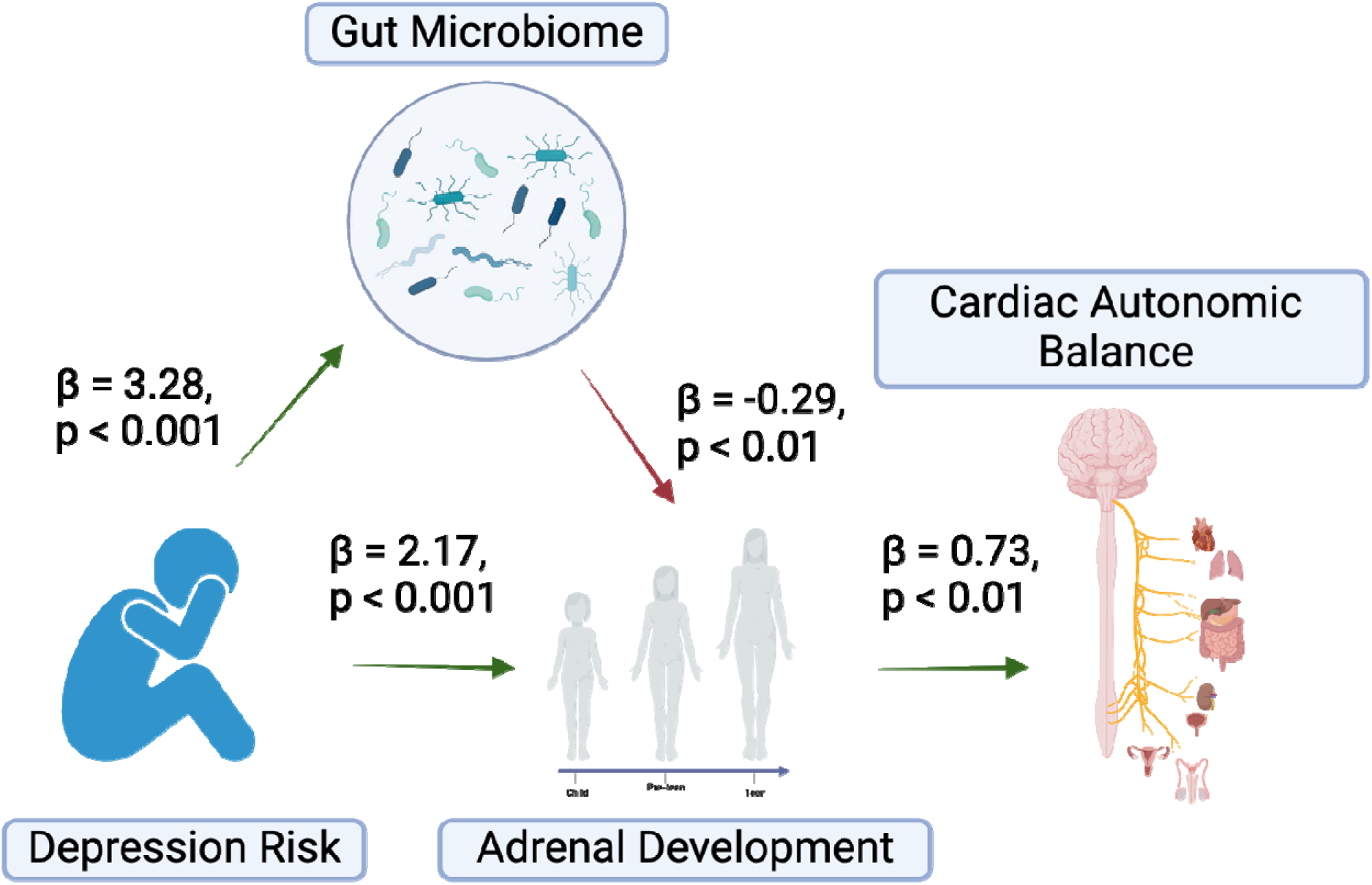
Gut Microbiota Mediates the Relationship Between Familial Depression Risk and Adrenal Development in Female Participants in Adolescence. We used lavaan to construct a mediation model with N=31 female participants to examine whether gut microbiota mediates the relationship between familial depression risk and adrenal development. The indirect effect of depression risk on pubertal development via gut microbiota was tested using the bootstrap method with 1000 resamples to obtain robust standard errors and confidence intervals. Our results showed that depression risk significantly predicted gut microbiota composition (β = 3.28, p < 0.001), which in turn significantly affected adrenal development (β = -0.29, p < 0.01). Additionally, the direct effect of depression risk on adrenal development was significant (β = 2.17, p < 0.001), and the mediated indirect effect through gut microb ota was also significant (β = -0.96, p < 0.05). Overall, these findings indicate a partial statistical mediation of gut microbiota in the link between depression risk and adrenal development. Finally, baseline cardiac autonomic balance was significantly associated with adrenal development.

**Figure 6.**
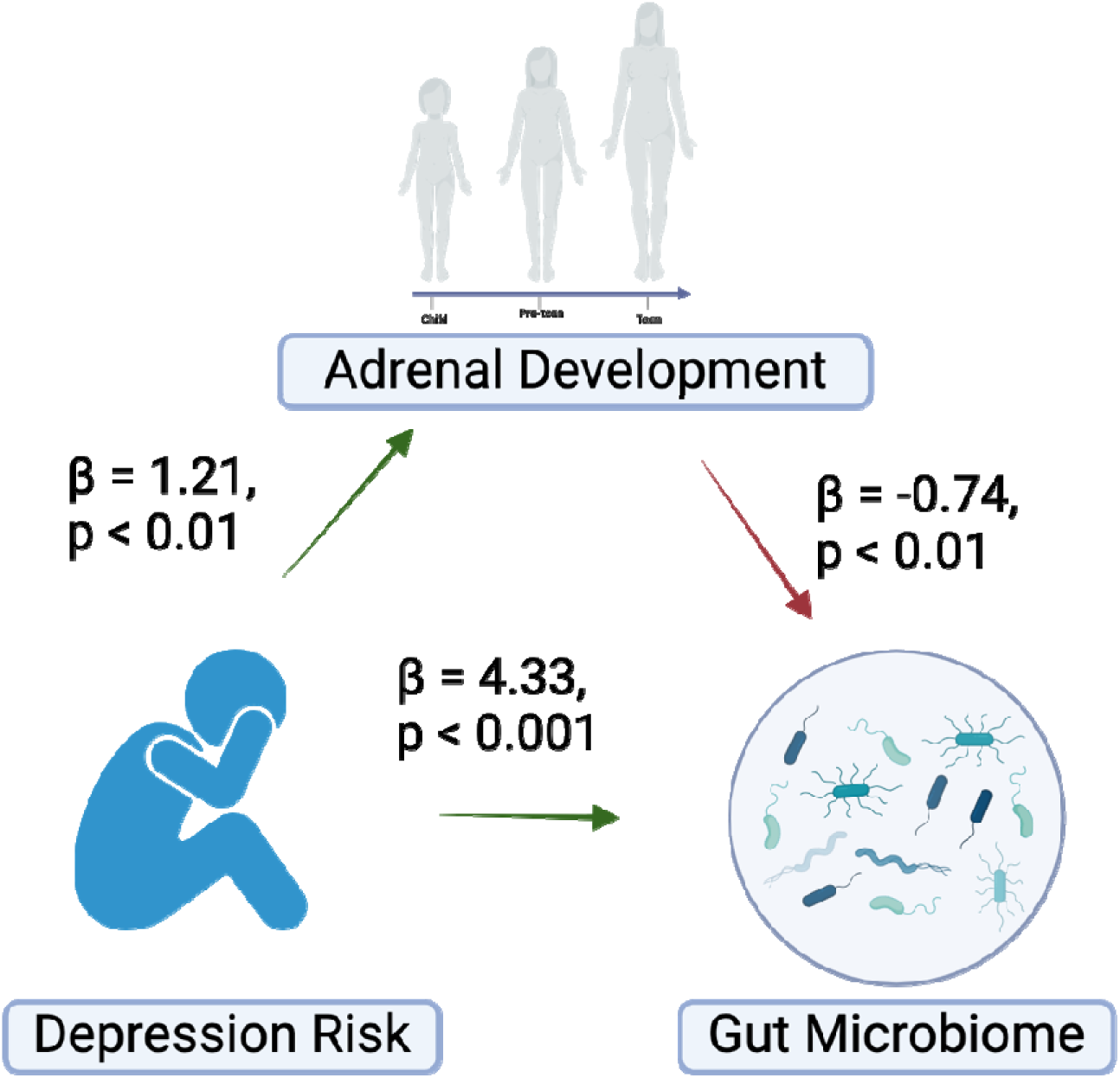
Adrenal Development Mediates the Relationship Between Familial Depression Risk and Gut Microbiota in Female Participants in Adolescence. We used lavaan to build a mediation model on N=31 female participants to test if adrenal development mediates the relationship between familial depression risk and gut microbiota. The indirect effect of depression risk on gut microbiota through adrenal development was tested using the bootstrap method with 1000 resamples to obtain robust standard errors and confidence intervals. Our results revealed that depression risk significantly predicted adrenal development (β = 4.33, p < 0.001), which in turn significantly influenced gut microbiota composition (β = -0.74, p < 0.01). Additionally, the direct effect of depression risk on gut microbiota was significant (β = 4.33, p < 0.001), and the indirect effect mediated by adrenal development was also significant (β = -0.90, p < 0.05). Overall, these findings suggest a partial mediation effect of adrenal development in the relationship between depression risk and gut microbiota.

Our results suggest that adrenarcheal development might significantly contribute to emerging mood vulnerability (Byrne et al., 2016; Ellis et al., 2019). However, our understanding of the mechanisms controlling the onset of adrenarche is limited (Rainey and Nakamura, 2008; Witchel et al., 2020). Adrenarche occurs before gonadarche and triggers the initial increase in adrenal androgens (DHEA/DHEA-S) that can influence immune function and microbial composition (Han et al., 2021; Prall and Muehlenbein, 2018; Rosenfield, 2021). The earlier appearance of these steroids in girls might prolong the period during which host-microbe-immune interactions can influence mood development (Liimatta et al., 2017; Rosenfield, 2021), offering one potential explanation for why adrenarche, but not gonadarche, had the strongest links to both *Prevotella* and depression risk in our study. The endocrine, microbial, and autonomic pathways connecting these observations are still not well understood, and further research is needed to identify the underlying mechanisms. Prospective studies that sample hormones, microbiota, and vagal activity before, during, and after adrenarche are now necessary to clarify causality and find modifiable targets for prevention.

The study has several limitations. First, our mediation analysis in females (n = 31) has limited power, and larger cohorts are needed for replication. Second, we lacked direct hormonal assays (such as DHEA, estradiol, testosterone, and cortisol), which would clarify the relationship between endocrine function and the microbiota. Third, dietary data were limited to whole-grain intake, preventing us from adjusting for fiber diversity or macronutrient profiles known to influence the relative abundance of *Prevotella*. Fourth, only 11% of participants met criteria for an anxiety disorder, which may indicate a bias toward lower comorbidity. Finally, indirect measures of adrenarche and vagal function, along with the cross-sectional design, restrict the ability to infer causality.

Nevertheless, the findings of the current study shed light on sex differences and depression risk during adolescence. Biological and lifestyle factors associated with increased familial depression risk may disrupt gut microbiota, adrenarcheal development, and stress reactivity. Understanding the mechanisms underlying adolescent depression could lead to new prevention and intervention strategies.

## Authors’ contributions

Conceptualization/Funding acquisition LB, CL, AM, BM; Data generation BM, AM; Data Curation – AK, EJ, LB; Formal Analysis – AK, EJ, LB; Original Draft – AK, EJ, LB; and Reviewing and Editing – AK, EJ, LB, CL, AM, BM.

## Ethics approval and Consent to participate

Written informed consent was obtained from all participants and their caregivers (parent consent and child assent).

## Acknowledgements.

The authors would like to thank The Center for Neural Basis of Cognition at the University of Pittsburgh and Carnegie Mellon University for supporting this work with the Community Collaboration Award.

